# Electrode position, distance, size, and orientation determine efficacy of cervical epidural stimulation to recruit forelimb muscles in rats

**DOI:** 10.1101/2025.09.05.674051

**Authors:** Andrés Pascual-Leone, Vishweshwar Tyagi, Ahmet S. Asan, Pedro E. Rocha-Flores, Ovidio Rodriguez-Lopez, Walter Voit, James R. McIntosh, Jason B. Carmel

**Affiliations:** Neurology, Columbia University, New York, NY; Neurological Surgery, Columbia University, New York, NY; Biomedical Engineering, University of Texas at Dallas, Richardson, TX; Center for Engineering Innovation, University of Texas at Dallas, Richardson, TX; Materials Science and Engineering, University of Texas at Dallas, Richardson, TX

**Keywords:** spinal cord stimulation, dorsal root entry zone, cervical epidural stimulation, motor threshold, evoked potential

## Abstract

Target engagement determines therapeutic efficacy. In spinal cord stimulation (SCS), efficacy depends on the attributes of the electrodes used to deliver stimulation, particularly their position, interelectrode distance, size, and current orientation. We tested the effects of these electrode parameters using cervical stimulation to elicit forelimb muscle responses in the rat. We implanted devices over the C6 dorsal root entry zone (DREZ) in eight rats and measured responses in six forelimb muscles. We used custom designed electrode arrays to systematically vary position, distance, and size with linear arrays, and current orientation with circular arrays. Stimulation consisted of biphasic and pseudomonophasic pulse shapes with bipolar or distant return, and a high-definition montage with four returns surrounding a central contact to increase local current density. Efficacy was calculated from the motor thresholds of recruitment curves, and parameters were compared using linear mixed models. For position, stimulation over the DREZ was most effective, reducing thresholds by 25.9% (*p* = 0.0002) relative to the midline; efficacy decreased as the electrode was positioned medial or lateral to the DREZ. For distance, placing the electrodes farther apart significantly improved efficacy (*p* = 0.0022). For size, large electrodes increased SCS efficacy, reducing thresholds by 21.5% (*p* = 0.0026) relative to small electrodes. After accounting for position and distance, current orientation did not affect SCS efficacy. Lastly, the high-definition montage decreased efficacy, increasing thresholds by 16.7% (*p* = 0.003) relative to monopolar stimulation with a distant return. In conclusion, electrode position, distance, and size had the greatest effect on SCS efficacy, with the optimal parameters combining large electrodes at the DREZ position with a distant return. These results have the potential to lower the current needed to engage cervical spinal circuitry and to inform the design of SCS systems to improve dexterity in people with neurological injury or disease.

## 1 Introduction

Therapeutic efficacy in neuromodulation depends on effective and selective engagement of the intended neural circuits. In spinal cord stimulation (SCS), targeted engagement is determined by a large space of electrical stimulation parameters, including electrode position, interelectrode distance, contact size, and current orientation. Although SCS is widely employed in both research and clinical care, electrode parameters remain largely unexplored, in part because of the difficulty of altering these parameters in a controlled manner. To address this gap, we evaluated epidural stimulation parameters in rats using custom built arrays to isolate their individual contributions to SCS efficacy. These devices were patterned using photolithography, enabling the electrodes to be systematically varied according to the parameters we tested [1–3].

Both animal and computational models of SCS indicate that dorsal epidural stimulation at low intensities primarily recruits large diameter afferent fibers [4–6]. These afferents synapse both directly and indirectly onto motoneurons and generate motor responses when stimulated at higher intensities [7, 8]. The resulting effect of afferent activation can be quantified using electromyography (EMG), which provides the ability to directly observe motor-evoked potentials (MEPs) as physiological response to stimulation. Previous studies have shown strong concordance between stimuli that produce consistent MEPs and those that promote adaptive plasticity [9, 10]. Experimental protocols can therefore be designed to compare stimulation parameters and optimize therapeutic efficacy of SCS using motor threshold as the primary outcome measure.

Electrode position relative to the spinal cord influences motor recruitment [4, 11– 14]. Prior studies have shown that lateral epidural stimulation is more effective than medial stimulation in the rat; the most effective electrode position is over the dorsal root entry zone (DREZ) [12, 13]. Similarly, in human intraoperative mapping experiments, epidural stimulation over the DREZ resulted in larger MEPs than midline stimulation in the cervical spinal cord [14]. In addition to the proximity of the electrodes to the dorsal root when positioned over the DREZ, the sharp change in the angle of afferent fibers as they enter the spinal cord may explain the enhanced stimulation efficacy observed near the DREZ [11]. Using arrays that spanned multiple degrees of laterality relative to the midline, we hypothesized that stimulation centered at the DREZ would be optimal, with a loss of SCS efficacy as electrodes were positioned laterally or medially from the DREZ.

Beyond the absolute position of the electrodes over the spinal cord, the interelectrode distance, or the spacing between the anode and cathode, determines the volume of tissue activated and the depth and spread of the resulting electric field. Computational studies in SCS have demonstrated that increasing the inter-electrode distance enhances neuron polarization up to a point [15]. However, if electrodes are placed too close to one another, the current preferentially shunts through the highly conductive cerebrospinal fluid rather than penetrating the higher resistance spinal cord tissue. Conversely, increasing the distance broadens and deepens the electric field, which can lower motor thresholds. Hence, we hypothesized that greater interelectrode distance would increase efficacy and result in lower motor threshold.

Modeling studies have demonstrated the impact of electrode contact size on neural activation efficacy [16, 17]. In clinical SCS studies for chronic pain, larger paddle lead devices allowed lower amplitudes with longer sustained pain relief compared to smaller percutaneous contacts [18, 19]. Similarly, in animal models, larger contacts activated target motor neurons more effectively than small contacts, while very small (100-200 µm) contacts in densely packed configurations might enable higher selectivity [20, 21]. We hypothesized that electrodes with a larger surface area would be more effective.

Stimulation waveform and electrical current direction also determine the efficacy of SCS. Contrary to cortical stimulation, monopolar monophasic cathode stimulation is more effective than anode stimulation for epidural spinal stimulation [22–24]. Alam et al. reported that monophasic bipolar stimulation across segments is superior to monopolar stimulation at a single segment for eliciting MEPs and subsequently showed that bipolar stimulation enhanced forelimb recovery after a C4 spinal cord injury in rats [24, 25]. Furthermore, computational models have demonstrated the importance of cathode positioning relative to the spinal cord [26, 27]. Cuellar et al. used a 4-contact array to stimulate at 45-degree radial increments, however, an optimum orientation could not be identified [28]. Computational models have shown that activation of nerve fibers by external stimulation is governed by changes in the electrical potential along the fiber [26, 29]. This suggested that stimulation waveform, polarity, and current orientation are likely important parameters determining SCS efficacy.

Precise testing of electrode position, interelectrode distance, contact size, and current orientation has been limited by a lack of technology to precisely pattern electrodes. For instance, the 4-contact array used by Cuellar et al. required combined activation of three contacts to select 45-degree increments of electrical current orientation, which is likely to produce complex current spread [28]. We and others have used photolithography to enable the printing of electrodes of any size or position [1, 3, 30–33]. We used a softening polymer that creates a tight neural interface with the underlying dura. We patterned these devices with electrodes specifically designed to test electrode position, interelectrode distance, contact size, and current orientation. We also tested biphasic and pseudomonophasic waveforms, and a high-definition montage that has been shown to increase local current density for brain stimulation [34, 35]. We have previously demonstrated that MEP recruitment curves allow accurate estimation of motor thresholds across multiple muscles [36]. We used custom electrode arrays and randomized stimulation to record MEP recruitment curves, from which motor thresholds were estimated. These thresholds were analyzed using linear mixed models to determine the efficacy of different electrode parameters and stimulation waveforms.

## 2 Methods

### 2.1 Summary

We stimulated the cervical spinal cord of eight rats using different electrode configurations while recording MEP responses from six muscles of the left forelimb (Fig. 1A). These muscles were chosen to represent different segments of the cervical enlargement. The effects of electrode position, interelectrode distance, electrode size, and current orientation on stimulation efficacy were tested using a linear array (Fig. 1B, left) with small and large contacts and a circular array (Fig. 1B, right). Figure 1C shows a computer rendering of an SCS device implanted through a C4 laminectomy. The substrate of the SCS device is a softening polymer; once placed on the spinal cord, the softening polymer allows the array to conform to the curvature of the underlying dura. The three experiments consisted of stimulating over the C6 segment (Fig. 1D1–D3) using the small and large contacts of the linear array and the circular array, with the center electrode positioned over the dorsal root entry zone (DREZ). Figure 1E shows the relative positions of the two arrays overlaid on one another, with overlapping contacts at the midline position. We used the recorded motor-evoked potentials to estimate motor thresholds from recruitment curves [36] across the six recorded muscles (Fig. 1F).

**Figure 1.**
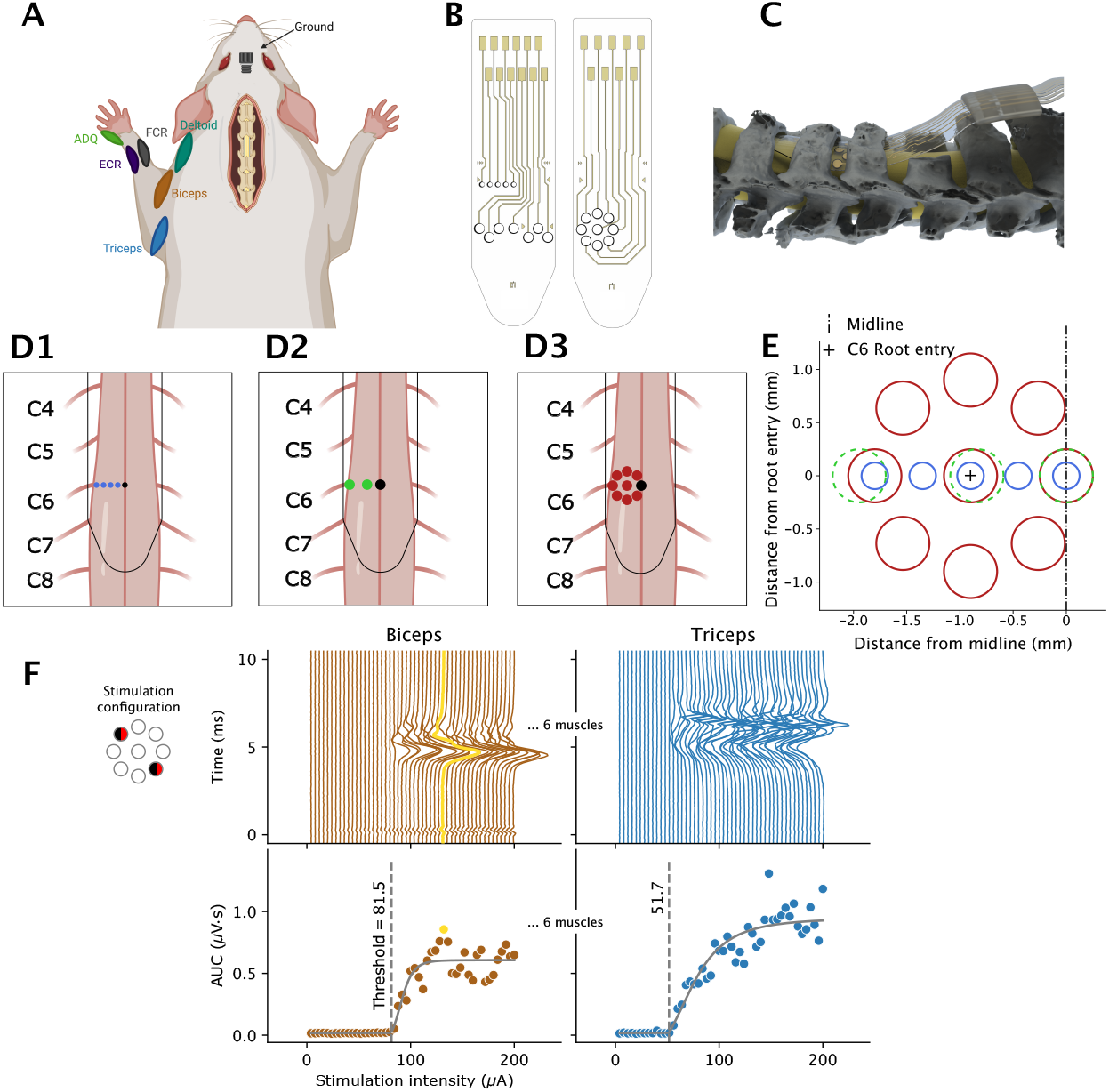
Experimental setup. **(A)** EMG electrodes implanted in six muscles of the rat’s left forelimb. A ground electrode was placed at the skull midline, over the olfactory bulbs and between the eyes. Created in BioRender. Pascual-Leone, A. (2025) https://BioRender.com/18tnpip **(B)** A linear array (left) consisting of small and large contacts and a circular array (right) were designed to evaluate different combinations of stimulation parameters. **(C)** Computer rendering based on micro-CT showing epidural insertion of an electrode following a C4 laminectomy. The electrode, fabricated from a softening polymer, conforms to the spinal cord curvature. **(D1–D3)** Electrodes were placed epidurally on the dorsal surface of the spinal cord, targeting the dorsal root entry zone at the C6 segment. Black circles mark the midline positions. D1: small contacts. D2: large contacts. D3: circular array. **(E)** Schematic overview of the spatial overlap across the three experiments, showing their relative alignment. **(F)** Example motor-evoked potentials (MEPs) recorded from the biceps and triceps in a rat when stimulation is applied in the root-perpendicular orientation. Top panels: motor-evoked potentials. Biphasic stimulation is represented by dual-shaded contacts, with red–black denoting an anode–cathode sequence and black–red a cathode–anode sequence. Bottom panels: MEPs are quantified by their area under the curve (AUC), and curves (solid lines) are fit to yield threshold estimates (vertical dashed line).

The mean threshold estimates across the six muscles (Fig. 2A1,A2) were used to analyze the efficacy of different stimulation parameters in reducing threshold using linear mixed models (Fig. 2B1,B2). This approach assumed that any bipolar biphasic stimulation can be decomposed into contributions from individual positions (Fig. 2B1,B2, orange bars), plus a distance effect between the positions (purple bars), and, in the case of large electrodes, an effect of large electrode size (green bar), and, in the case of the circular array, an effect of orientation (see Methods 2.8).

**Figure 2.**
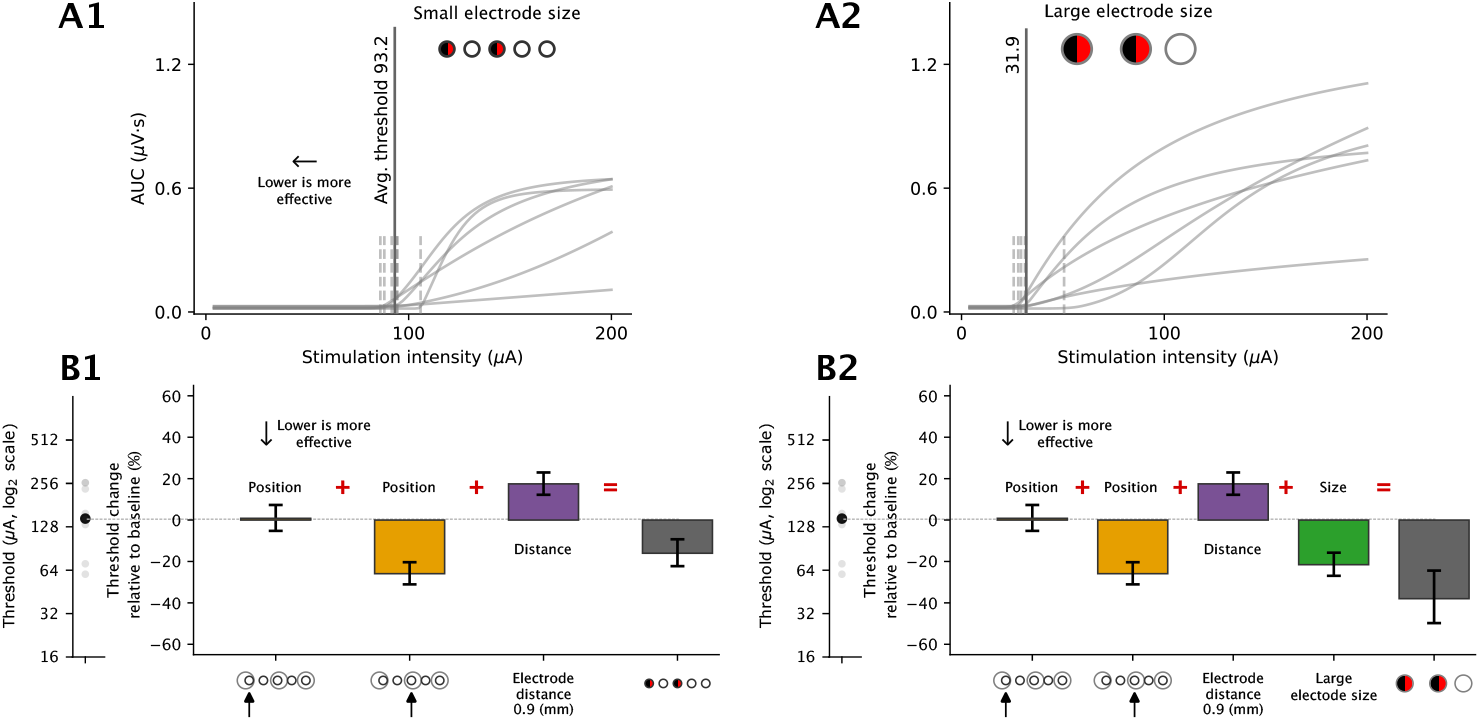
Linear mixed model decomposition. **(A1)** Example recruitment curves for six forelimb muscles in a rat under bipolar biphasic stimulation between the center and far lateral positions using small electrodes. Gray solid curves represent recruitment curves of individual muscles, gray vertical dashed lines denote their thresholds, and a gray vertical line denotes the mean threshold across muscles. **(A2)** Same as (A1) but with large electrodes, showing lower thresholds across muscles. **(B1)** Decomposition of the mean threshold for bipolar stimulation into additive contributions from the center and far lateral positions, along with an additional effect of interelectrode distance. Effects are expressed as percent change in threshold relative to baseline monopolar stimulation at the midline with small electrodes. **(B2)** Same as (B1), with an additional contribution from large electrode size, which further reduces the threshold.

### 2.2 Spinal cord electrode array design and fabrication

Spinal cord electrode arrays (∼100 *µ*m total thickness) were fabricated on a softening thiol-ene/acrylate polymer substrate using standard photolithographic microfabrication methods [1, 3, 33]. The device architecture consisted of gold interconnects (300 nm) with an electrode coating layer (titanium nitride), encapsulated between two polymer layers and patterned using lithography, wet etching, and reactive-ion etching processes before water release. Two sizes of electrodes were patterned: large electrodes (diameter: 500 *µ*m) and small electrodes (diameter: 250 *µ*m). Device stimulation properties were characterized using standard electrochemical characterization techniques [37] to ensure similar charge injection capacities.

### 2.3 Surgical procedures

Eight adult female Sprague Dawley rats were used in this study for a terminal physiology experiment. All procedures were conducted in compliance with the guidelines of the Institutional Animal Care and Use Committee at Columbia University in New York, NY, and followed aseptic techniques during a terminal physiology experiment. Experiments were performed under anesthesia to enable testing with different devices.

Surgical procedures followed our published protocols [10, 12] and are detailed in the Supplementary Methods. Key details are summarized below. EMG was recorded from eight different muscles: left extensor carpi radialis (ECR), flexor carpi radialis (FCR), biceps, triceps, abductor digiti quinti (ADQ), deltoid, biceps femoris, and right biceps. A laminectomy was performed to expose the C4 cervical spinal cord. Arrays were then carefully inserted into the dorsal epidural space with electrodes positioned over the C6 root. We used bony and anatomical landmarks to identify the root entry zone. The intervertebral foramen also provided an opening for visual confirmation of the target site. As various SCS arrays were used in this study (Fig. 1D1–D3), they were replaced sequentially, starting with the large linear electrodes, followed by the small electrodes, and then the circular array. When using the circular array, the center electrode was placed over the C6 DREZ. When using the linear electrode configuration, the array was advanced segmentally to target each DREZ at C5 and C6 with both the large and small contact rows. Thus, both contact sizes stimulated both levels.

### 2.4 Electrophysiology and stimulation

Connectors for the spinal cord array and EMG wires were attached to a headstage ZIF-connector (Tucker-Davis Technologies; Florida, USA). The headstage connectors were used to interface the implanted electrodes with an amplifier (Tucker-Davis Technologies, PZ5), which was then connected to a real-time signal processing system (RZ2). A 16-channel IZ2H/Subject Interface constant current stimulator (Tucker-Davis Technologies) was controlled via a custom Matlab (R2022a) codebase. Single-pulse stimulation was delivered with various pulse shapes (biphasic and pseudomonophasic) with equal pulse widths of 200 *µ*s in their leading phase every 2 seconds. A charge balancing period of 800 *µ*s was used for pseudomonophasic pulses. In order to determine the maximum stimulation intensity for each rat and device, we manually increased intensity while monitoring the animal. The maximum was defined as the lowest intensity that first evoked visible trunk movement. To generate recruitment curves, stimulation intensities were subsequently linearly increased from 0 to the maximum intensity (350 ± 207 *µ*A) across 51 steps. This resulted in a step size of 6.43 ± 3.4 *µ*A. Stimulation patterns were randomly applied across all combinations of contacts for an individual array to control for order effects, excluding high impedance contacts from testing. EMG signals were sampled at 10 kHz and high-pass filtered using a 20 Hz cutoff 10th order IIR filter. A total of 155,484 MEPs across all rats and stimulation experiments were quantified from biceps, triceps, ECR, FCR, APB and ADQ ipsilateral to the side of stimulation. Contralateral biceps and biceps femoris were excluded from subsequent analyses as we were primarily interested in ipsilateral forelimb activation.

### 2.5 Power

We used differences in log_2_-transformed thresholds from 13 human participants [14] comparing midline and lateral epidural cervical stimulation in the triceps muscle (mean = 0.98, SD = 0.48). We planned a two-sided paired *t*-test on the differences in log_2_-transformed thresholds and controlled the familywise Type I error rate at *α* = 0.05 across a maximum of nine planned comparisons (the number of positions that can be compared on the circular array) using a Bonferroni correction. Under these assumptions, the required sample size to achieve 90% power was *n* = 8 rats.

### 2.6 Outcome measure

We defined the mean threshold across the six recorded muscles as our outcome measure. Thresholds were estimated simultaneously for all six muscles from recruitment curves (Methods 2.8). Electrode parameters that result in lower thresholds correspond to higher efficacy.

### 2.7 MEP size calculation

MEPs were quantified using area under the curve (AUC). In all experiments, EMG AUC was measured in a window between 1.5 ms and 10 ms after the spinal stimulation trigger. This window was chosen to capture activation of Type-I sensory afferents as well as direct activation of motor efferents and agrees with prior work in the rat cervical spinal cord [38] covering early and middle responses.

### 2.8 Estimation of recruitment curves and efficacy analysis

Throughout this manuscript, all logarithms are taken to base 2. We fit recruitment curves to MEP size data for each muscle to estimate thresholds. Let *a*^*r,c,m*^ denote the threshold for rat *r*, stimulation configuration *c*, and muscle *m*. We model thresholds on the log scale to assess multiplicative changes between stimulation configurations. On this scale, a one unit change represents a twofold change in threshold on the original scale. We define the mean threshold *a*^*r,c*^ as the arithmetic mean of the log transformed thresholds log(*a*^*r,c,m*^) across the six muscles (Eq. 1), which is also equivalent to the log transformed geometric mean of thresholds on the original scale (Eq. 2). In the case of the linear array, which was tested at two cervical segments (C5 and C6), we additionally averaged the log transformed thresholds across segments.

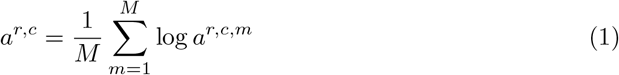

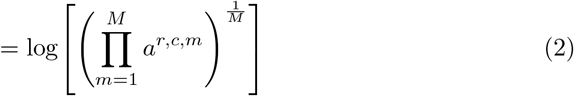

#### 2.8.1 Electrode position, interelectrode distance, and electrode size in linear array

Let 𝕃 = {LL, L, C, LM, M} denote the set of electrode positions in the linear array, labeled by their mediolateral positions left to right—far lateral (LL), lateral (L), center (C), lateromedial (LM), and midline (M). The configurations tested were either bipolar biphasic (*p*_1_, *p*_2_) ∈ 𝕃 × 𝕃, *p*_1_ ≠ *p*_2_, or monopolar biphasic (·, *p*) with *p* ∈ 𝕃. Both small and large electrode sizes were tested in this array. However, large electrodes were only available at *p* ∈ {LL, C, M}. For each configuration *c*, define position indicators

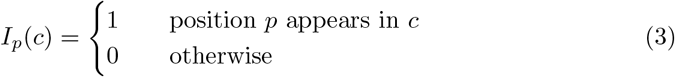

Also, define the size indicator

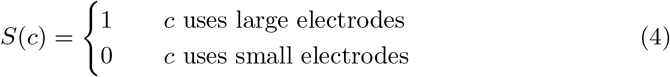

Let *d*(*p*_1_, *p*_2_) ∈ [0, *∞*) denote the Euclidean distance between positions *p*_1_ and *p*_2_. We define the distance term as,

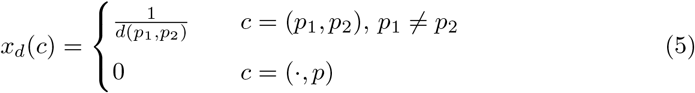

Then, the mixed model is given by

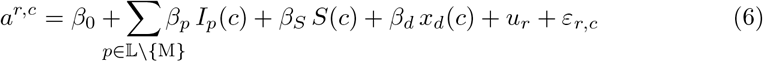

where *u*_*r*_ is a random intercept for rat 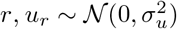, and *ε*_*r,c*_ ∼ 𝒩 (0, *σ*^2^). The intercept *β*_0_ corresponds to the monopolar (·, M) with small electrode size. Supplementary Table S1 shows the estimated fixed effects using this model.

#### 2.8.2 Electrode position, interelectrode distance, and current orientation in circular array

Let ℂ = {C, NE, N, NW, W, SW, S, SE, E} denote the set of electrode positions in the circular array, labeled by their cardinal positions and ordered in the anticlockwise direction, starting with the center. The circular array consisted of large electrodes, and the configurations tested were either bipolar biphasic (*p*_1_, *p*_2_) ∈ ℂ × ℂ, *p*_1 ≠_ *p*_2_, or monopolar biphasic (·, *p*) with *p* ∈ ℂ.

We use the position indicators and distance term from before (Eq. 3, 5). Additionally, let *θ*(*p*_1_, *p*_2_) ∈ [0, *π*) denote the axis orientation of bipolar contacts (*p*_1_, *p*_2_) relative to the E–W axis, i.e., *θ*(E, W) = 0 and increasing in the anticlockwise direction. We define 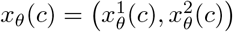 as,

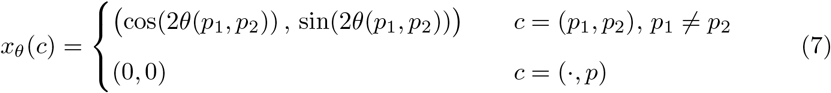

Then, the mixed model is given by

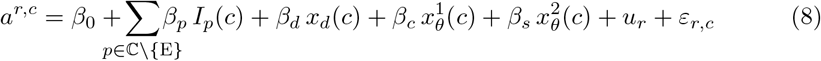

where *u*_*r*_ and *ε*_*r,c*_ are as in Eq. 6. The intercept *β*_0_ corresponds to the monopolar (·, E). Supplementary Table S2 shows the estimated fixed effects using this model.

In Eq. 7, the angle *θ*(*p*_1_, *p*_2_) is multiplied by 2 to account for axial orientation rather than direction. We have *C* cos(2*θ* − *ϕ*) = cos(2*θ*) (*C* cos *ϕ*)+sin(2*θ*) (*C* sin *ϕ*), hence, the overall amplitude *C* and phase offset *ϕ* of the orientation effect are derived from the fitted linear coefficients *β*_*c*_ and *β*_*s*_ corresponding to cos(2*θ*) and sin(2*θ*), respectively, by solving *β*_*c*_ = *C* cos *ϕ* and *β*_*s*_ = *C* sin *ϕ*. To determine the statistical significance of the overall orientation effect amplitude *C*, we performed a joint Wald test of the null hypothesis *H*_0_ : *β*_*c*_ = *β*_*s*_ = 0.

#### 2.8.3 Waveform, polarity, and high-definition return in circular array

Thus far, the configurations tested consisted of biphasic stimulation, hence we assumed (*p*_1_, *p*_2_) and (*p*_2_, *p*_1_) as equivalent. Using the circular array, we additionally verified this and evaluated the effects of polarity, waveform and return configuration at the center electrode. The eight tested configurations comprised four waveform and polarity combinations under a distant return and the same four combinations tested with a high-definition montage.

Let ℙ = {−, +} denote polarity at the center electrode, where − indicates cathode at center and + indicates anode at center. Let 𝕎 = BIP, PSM denote wave-form, indicating biphasic (BIP) and pseudomonophasic (PSM) pulse shape. For each configuration *c*, define indicators

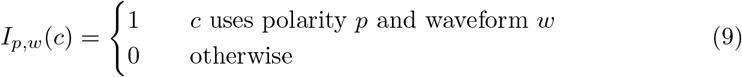

Define an indicator for the high-definition return,

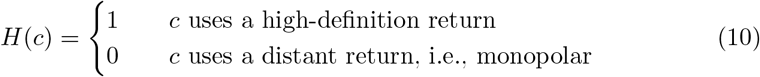

The mixed model is given by

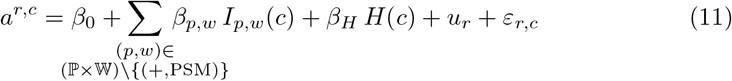

where 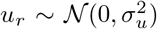 and *ε*_*r,c*_ ∼ 𝒩 (0, *σ*^2^) as in Eq. 6. The intercept *β*_0_ corresponds to pseudomonophasic pulse with anode at the center and cathode at a distant return. Supplementary Table S3 shows the estimated fixed effects using this model.

### 2.9 Statistics and reproducibility

All recruitment curves were fit using the python package hbMEP [36], and posterior median values were used as point estimates for motor threshold. The linear mixed models were fit using statsmodels [39]. All *p*-values in results are reported after correcting for multiple comparisons using the Holm–Bonferroni method [40]. All intervals reported are 95% confidence intervals and are not adjusted for multiple comparisons. The code to reproduce the results is available on GitHub (see Code availability).

## 3 Results

### 3.1 Position: Stimulation over the DREZ is most effective

The linear array consisted of five mediolateral positions (Fig. 1E), labeled from left to right—far lateral (LL), lateral (L), center (C), lateromedial (LM), and midline (M). The center (C) position was anatomically centered over the dorsal root entry zone (DREZ). Figure 3A,B show the motor thresholds estimated across eight rats for all configurations tested in this array, which included monopolar and bipolar biphasic stimulation. Both small and large electrodes were tested, however, large electrodes were available only at LL, C, and M. Figure 3C shows the fixed effect estimates from the linear mixed model (Eq. 6), expressed as percent change in threshold relative to the baseline configuration, chosen as monopolar stimulation at midline (M) using small electrodes. Negative values indicate lower thresholds and, therefore, greater efficacy.

**Figure 3.**
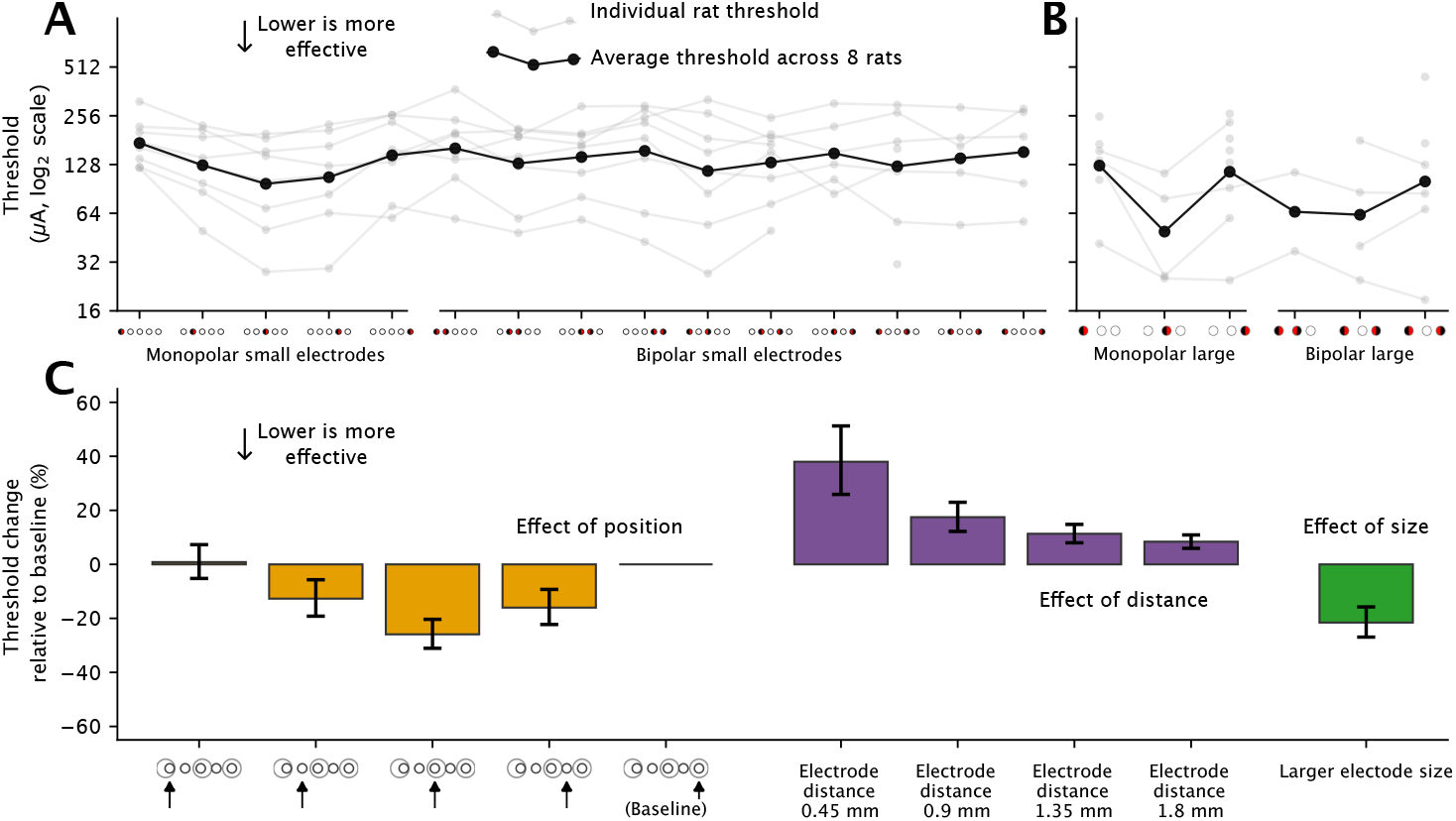
Effect of mediolateral position, interelectrode distance, and size in the linear array. **(A)** Per-rat (gray) motor thresholds and the average across rats (black) for monopolar and bipolar stimulation with small electrodes. **(B)** Same as (A) for large electrodes. **(C)** Mixed model estimates showing the effects of position, interelectrode distance, and electrode size, expressed as percent change in threshold relative to the baseline (monopolar stimulation at the midline using small electrodes). Lower values indicate greater efficacy. Center position over the DREZ, greater interelectrode distance, and larger electrode size are more effective and result in lower thresholds. Error bars represent mean ± SEM.

Electrode position significantly influenced efficacy, with stimulation over the DREZ producing the lowest thresholds (Fig. 3C yellow bars). Relative to the midline reference (M), the center position (C) was associated with the largest reduction of 25.9% (CI : (−35.7%, −14.6%), *p* = 0.0002). The adjacent lateromedial position (LM) showed a marginal decrease of 16%, which was not significant (CI : (−27.8%, −2.3%), *p* = 0.07), and was comparable to the adjacent lateral position (L). The far lateral position (LL) was comparable to the midline.

### 3.2 Distance: Electrodes farther apart are more effective

Increasing interelectrode distance was associated with lower thresholds, indicating increased efficacy at greater separation, with monopolar stimulation being the most effective (Fig. 3C purple bars). The effect of distance was significant (*p* = 0.0022) (Methods 2.8.1, Eq. 5,6), corresponding to a threshold increase of 38% at the closest tested separation of 0.45 mm (CI : (15.3%, 65.2%)) and a marginal increase of 8.4% at 1.8 mm separation (CI : (3.6%, 13.4%)) relative to monopolar stimulation.

### 3.3 Size: Larger contacts are more effective

Electrode size significantly affected efficacy (Fig. 3C green bar). Large electrodes yielded lower thresholds compared to small electrodes, corresponding to a reduction of 21.5% (CI : (−31.8%, −9.8%), *p* = 0.0026) relative to baseline.

Overall, SCS efficacy in the linear array was jointly determined by electrode position, interelectrode spacing, and electrode size. Placement over the DREZ, greater electrode spacing, and larger electrodes were associated with lower motor thresholds and greater efficacy.

### 3.4 Circular array shows consistent position and distance effect

The circular array was designed to assess whether current orientation influences stimulation efficacy. Since it was constructed on the same coordinate system as the linear array, comparable mediolateral positions were tested (Fig. 1E). The circular array consisted of eight cardinal positions and a center. In particular, the center (C), north (N), and south (S) positions were aligned over the DREZ, corresponding anatomically to the center (C) position in the linear array, while the east (E) position corresponded to the midline (M) of the linear array.

Figure 4A–C show the thresholds estimated across eight rats for all bipolar and monopolar configurations. Figure 4D shows the fixed effects of electrode position, distance, and orientation from the linear mixed model (Eq. 8), expressed as percent change in threshold relative to the baseline configuration, chosen as monopolar stimulation at E. Consistent with the linear array, both electrode position and distance significantly influenced efficacy (Fig. 4D, yellow and purple bars).

**Figure 4.**
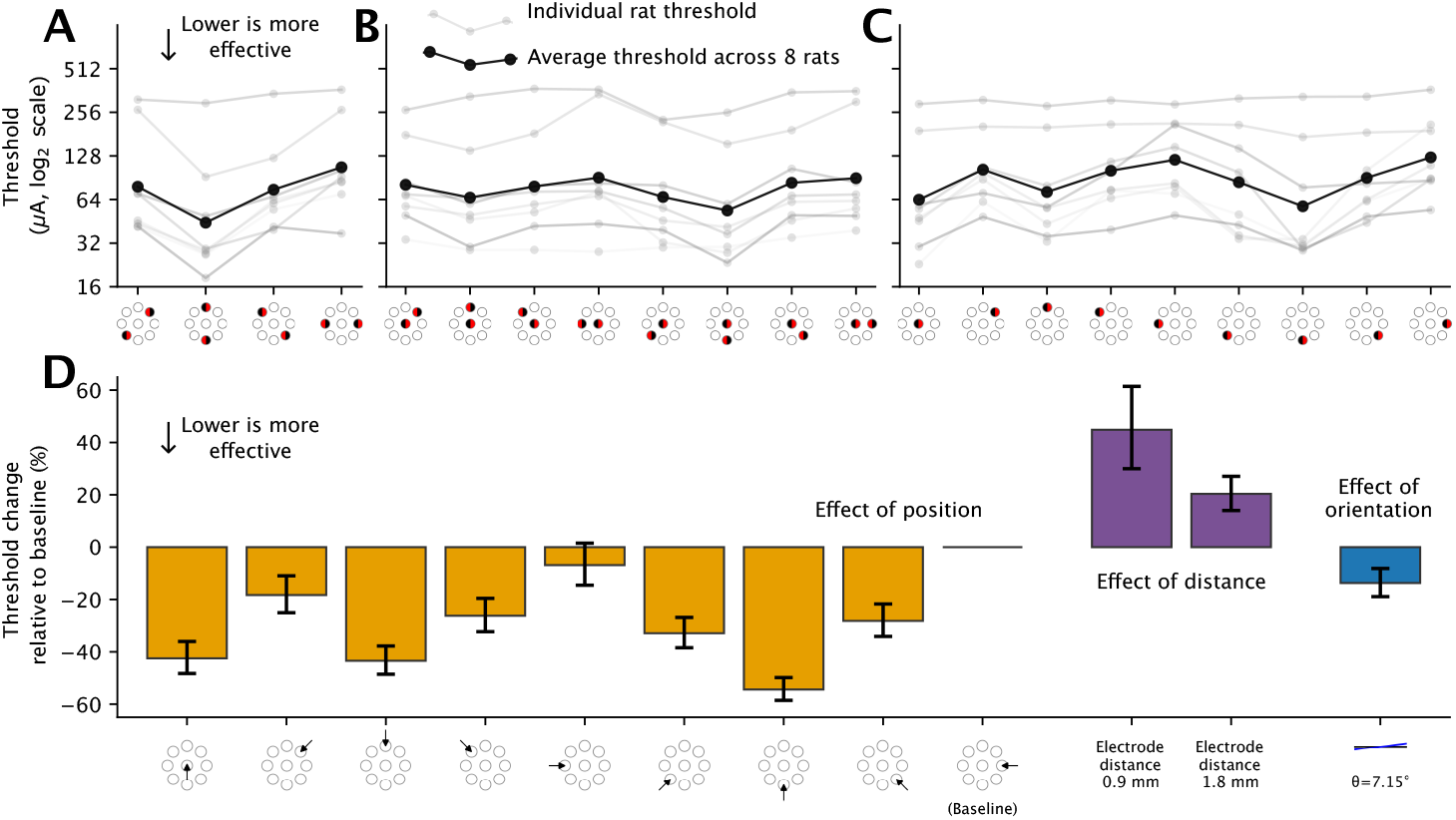
Effect of mediolateral and rostro-caudal position, electrode distance, and orientation in the circular array. **(A)** Per-rat (gray) motor thresholds and the average across rats (black) for diametrically opposite bipolar stimulation. **(B)** Same as (A) for radial bipolar stimulation. **(C)** Same as (A) for monopolar stimulation. **(D)** Mixed model estimates showing the effects of mediolateral and rostro-caudal position, interelectrode distance, and orientation, expressed as percent change in threshold relative to the baseline (monopolar stimulation at E). Lower values indicate greater efficacy. Positions aligned with the DREZ and electrodes farther apart are more effective, and orientation has minimal independent effect on efficacy. Error bars represent mean ± SEM.

Positions aligned over the DREZ produced the largest reductions in threshold relative to baseline. The south (S) position was the most effective, reducing thresholds by 54.4% (CI : (−62.2%, −45%), *p* = 1.7 × 10^−15^), followed by a 43.4% reduction at north (N) (CI : (−53.1%, −31.8%), *p* = 2 × 10^−8^) and a 42.5% reduction at center (CI : (−53.3%, −29.2%), *p* = 1.5 × 10^−6^). Adjacent positions (SW, SE, NW) also reduced thresholds significantly, whereas NE and the far lateral position (W) did not differ from baseline. Overall, efficacy increased as the electrodes were positioned closer to the DREZ.

Consistent with the linear array, interelectrode distance also significantly affected efficacy, with electrodes farther apart being more effective (*p* = 0.0025) (Fig. 4D purple bars). Closely spaced radial bipolar stimulation (0.9 mm) corresponded to a threshold increase of 44.9% (CI : (17.1%, 79.1%)), and diametrically opposite stimulation with greater separation (1.8 mm) corresponded to an increase of 20.4% (CI : (8.2%, 33.8%)) relative to monopolar stimulation.

### 3.5 Orientation has minimal independent effect

After accounting for electrode position and interelectrode distance, orientation contributed minimally to efficacy (Fig. 4D blue bar). The orientation term was not significant using a Wald test (*p* = 0.12, Methods 2.8.2), and the estimated effect across orientations was small relative to the effects of position and distance. Thus, once mediolateral and rostro-caudal placement and interelectrode spacing were controlled for, there was no evidence of a significant orientation effect on threshold.

### 3.6 Polarity: Cathode stimulation is more effective

To determine whether waveform polarity influences motor recruitment, we tested pseudomonophasic and biphasic pulse shapes using the circular array (Fig. 5A). Additionally, to assess whether increasing local current density around the DREZ affects motor threshold, we tested a high-definition montage [35], consisting of a central electrode (C) surrounded by four adjacent electrodes (NE, NW, SE, SW) of opposite polarity. Figure 5B shows the fixed effects of waveform, polarity and the high-definition montage, expressed as percent change in threshold relative to the baseline configuration, chosen as pseudomonophasic anode stimulation at center.

**Figure 5.**
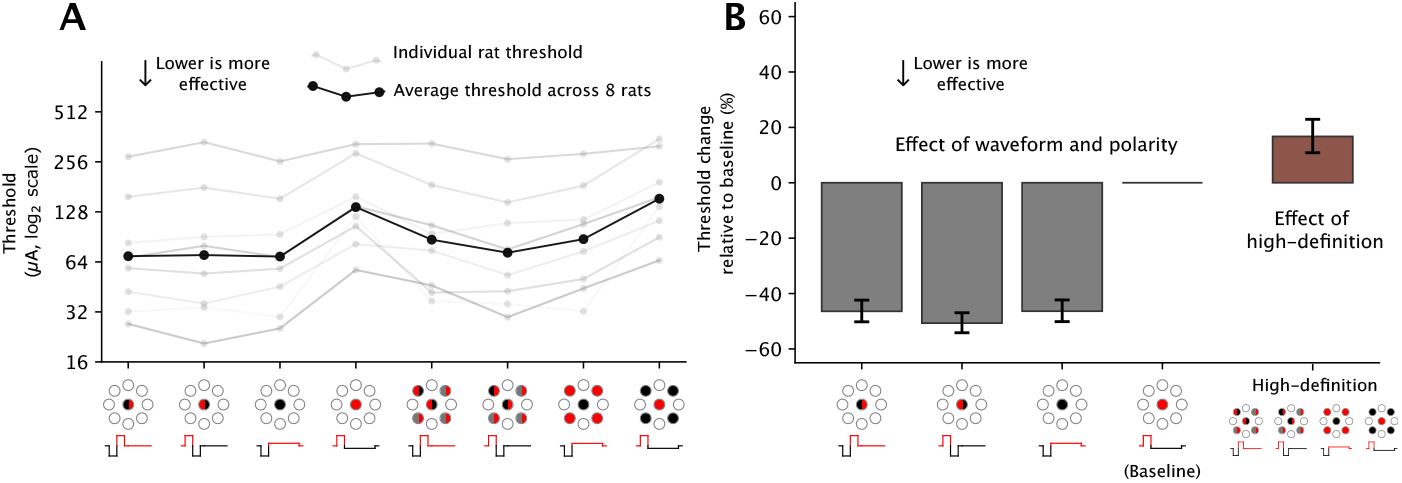
Effect of waveform, polarity, and high-definition return in circular array. **(A)** Per-rat (gray) motor thresholds and the average across rats (black) for biphasic and pseudomonophasic pulse shapes, cathode-first and anode-first polarity, and the high-definition return. **(B)** Mixed model estimates showing the effects of waveform, polarity, and high-definition return, expressed as percent change in threshold relative to the baseline (pseudomonophasic anode stimulation at the center). Lower values indicate greater efficacy. Cathode stimulation reduces thresholds relative to anode stimulation, while the high-definition return increases thresholds compared to a distant return. Error bars represent mean ± SEM.

Compared to the baseline, pseudomonophasic cathode stimulation reduced motor thresholds by 46.4% (CI : (−53.5%, −38.1%), *p* = 3.7 × 10^−17^). This was comparable to biphasic stimulation, both for cathode-first pulse, which reduced thresholds by 46.4% (CI : (−53.6%, −38.2%), *p* = 3.7 × 10^−17^), and for anode-first pulse, which reduced thresholds by 50.7% (CI : (−57.3%, −43.1%), *p* = 1.4 × 10^−21^). This indicates that cathode stimulation at the DREZ is significantly more effective than anode stimulation, and comparable to biphasic stimulation.

### 3.7 High-definition decreases efficacy over distant return

After controlling for waveform polarity, the high-definition montage resulted in significantly higher thresholds compared to monopolar stimulation with a distant return (Fig. 5B brown bar), corresponding to a 16.7% increase in threshold (CI : (5.5%, 29.1%), *p* = 0.003).

## 4 Discussion

In this study, we aimed to identify the electrode parameters that influence SCS efficacy when targeting forelimb motor recruitment in the rat. We used custom-built arrays to systematically vary electrode position, interelectrode distance, electrode size, current orientation, pulse shape, and polarity. Because these parameters cannot be varied in isolation, we used linear mixed models to estimate their individual contributions to SCS efficacy, resulting in five key observations. First, electrode position over the DREZ was most effective, reducing motor thresholds by 25.9% relative to the midline, with efficacy decreasing as electrodes were positioned more medially or laterally. Second, interelectrode distance significantly affected efficacy, with electrodes farther apart being more effective and producing lower thresholds. Third, larger electrodes increased efficacy, reducing thresholds by 21.5% relative to small electrodes. Fourth, cathode stimulation at the DREZ was more effective, reducing thresholds by 46.4% relative to anode stimulation. Finally, after accounting for electrode position, distance, and size, orientation did not have a significant effect on SCS efficacy.

For electrode position, stimulation over the DREZ resulted in lower motor thresholds compared to more medial and lateral positions. This is consistent with our previous work and that of others showing that targeting the DREZ in the cervical spinal cord with dorsal epidural stimulation yields larger MEPs [6, 12, 14]. In a finite element model of epidural stimulation of the lumbar spine, Rattay et al. identified anatomical hotspots at regions of sharp curvature just before entry into the spinal cord and at the point of entry itself, i.e., the DREZ [4]. These findings suggest that local changes in fiber orientation at the DREZ may provide a favorable environment for electrical stimulation, whereas more medial or lateral locations lack such changes and require higher stimulation intensities to evoke comparable responses.

In addition to electrode position, we also evaluated the effect of interelectrode distance, or the spacing between the anode and the cathode. We found that this effect was significant, with electrodes placed farther apart being more effective. In finite element modeling of transcranial direct current stimulation, previous work has suggested that increased electrode separation reduces current shunting through the scalp and CSF [41]. Potentially, electrodes in close proximity result in a greater degree of CSF shunting, leading to a relative loss of neuronal activation and increasing the required stimulus intensity.

For electrode size, larger electrodes (500 *µ*m in diameter) decreased the threshold compared to smaller electrodes (250 *µ*m) when electrode position and distance were held constant. Electrode size may not significantly affect the magnitude of the electrical field generated, but larger electrode contacts improve field uniformity [42], which in turn may influence the activated neural structures. Further studies are needed to determine whether there is a trade-off between efficacy, which favors large electrodes, and selectivity of the muscles innervated at the stimulated level, which may be greater with smaller electrodes.

Orientation had no significant effect on spinal cord stimulation efficacy, unlike brain stimulation when the effects of location and electrode distance are fixed. Electrical current vector-dependent motor responses have been extensively demonstrated in cortical stimulation studies [43–46]. In the spinal cord, computational models have demonstrated the importance of direction of the electrical field relative to the underlying axonal fibers [22, 23, 47]. We predicted that electrical current aligned to the root entry would be most effective, however, when controlling for position and inter-electrode distance, we found that the vector of stimulation minimally impacted MEP threshold alone. This potentially speaks to a relatively larger influence on efficacy by the proximity of contacts to the DREZ rather than to the alignment of the dorsal fibers that target it. This is likely influenced by the electrical field orientation relative to the underlying neural elements [5, 26, 29] and the properties of the anatomical structure [26, 48]. Previous groups have lacked the contact density in arrays to comprehensively assess the impact of electrode orientation. Our circular array contained contacts designed to align parallel and perpendicular to both the spinal cord and the dorsal roots. We found that an electrical current vector 7.15 degrees anticlock-wise to midline-lateral yields the most effective stimulation, but to a marginal and non-significant change.

Likewise, a high density montage, modeled off of cortical stimulation, decreased spinal cord stimulation efficacy, rather than increasing the efficacy of as seen in brain stimulation. Our circular array allowed for high-density configurations, a central cathode surrounded by multiple anodes, that provided increased current density over the DREZ. However, we found this lead to decreased efficacy in comparison to a monopolar stimulating electrode with a distant return. This may in part be explained similarly to interelectrode distance where the high-definition configuration likely led to greater current shunting than expected. Potentially, this configuration leads to greater selectivity (isolation of a singular target muscle) compared to a more broad stimulus, but future studies will have to assess the trade-off between efficacy and selectivity.

This study utilized methods novel to the field of neurophysiology that allowed the testing of parameters not previously investigated in epidural spinal cord stimulation. Our group designed and built electrode arrays with the direct purpose of testing specific electrode parameters, like current orientation and a high-definition array with the circular electrode array. This question-driven electrode array creation paired with animal studies allowed for rapid and comprehensive investigation of a variety of stimulation parameters, which was enabled by leveraging the recruitment curve analysis methods we have previously developed. We extended this analytical framework to incorporate mixed effect modeling to assess the relative contribution of each parameter and directly compare isolated effects on efficacy. This is a key difference from prior literature where effects of some of the parameters tested (position, orientation, etc.) have been studied, but whose individual effects have been confounded by other parameters. We also tested a novel high-definition configuration to mimic that of previously studied high-definition arrays in cortical stimulation which did not yield increased efficacy contrary to brain stimulation outcomes.

There are a few limitations of this study worth noting. First, while motor threshold is a well established measure of motor system activation, improving function is likely a far more complex process. Some comparisons, especially of the large linear electrodes, were limited by the number of rats with successful recruitment curves. This may be due to the array shifting during the experiment or being slightly off center during initial placement leaving some contacts only partially in contact with the spinal cord. However, a clear relationship was observed where larger size increased efficacy of stimulation. In addition, our customized arrays relied on anatomical landmarks for appropriate positioning of the electrode without radiographic confirmation. We are currently generating new arrays for subsequent experiments that contain radiopaque fiducials to confirm proper placement with x-ray. This will be particularly important for experiments conducted in awake rats. Although the current studies were performed in anesthetized rats, previous studies showed very similar effects of spinal cord stimulation in rats anesthetized with ketamine and xylazine compared with awake rats [10, 12]. Our electrodes were designed to test parameters within a single spinal segment. The most activated muscles would likely differ if targeting the DREZ of segments either above or below C6, as shown by other investigators [6]. Future studies should examine the interaction of stimulation across segments, similar to what we have done in mapping the human spinal cord [14].

Given a single combination of parameters is unlikely to be universally optimal, future studies should investigate how these parameters interact to better understand their combinatorial effects. Additionally, some parameters are fundamentally linked. For example, it is not possible to change orientation of current flow without simultaneously changing the position of the active electrodes. Advances in implantable technologies could improve both the delivery of electrical current and measurement of the effects through sensing electrodes. Measurement of spinal cord extracellular compound action potentials could serve as real-time physiological biomarkers to guide stimulation parameter selection. This approach may ultimately enable tailored stimulation strategies to support functional recovery, such as finding optimal stimulation paradigms for recovery of reaching or grasping movements in patients with loss of arm and hand function. Given the immensity of the parameter space available for exploration and optimization in epidural spinal cord stimulation, the use of rodents to advance the field is essential. While there are notable differences between rats and humans in motor pathways, the neural structures relevant for dorsal epidural stimulation, primarily the large diameter afferents, are largely conserved between the two species. Therefore, the use of rodents can provide conclusions to specific stimulation questions impossible to test in humans that can later inform clinical SCS electrode design.

This study demonstrates the potential for customized electrodes in combination with mixed model analyses to improve our understanding of dorsal epidural stimulation through targeted experiments with arrays designed to assess specific questions and isolate specific stimulation parameter effects. We investigated the relative impact of polarity, inter-electrode distance, medio-lateral position, contact size, current orientation, and high-definition configurations on stimulation efficacy. This experimental design allows for future targeted experiments in both animal and clinical trials aimed at improving motor recovery through optimization of epidural spinal cord stimulation. The continued advancement in stimulation efficacy significantly could improve clinical utility of stimulation. Higher efficacy would yield similar effects of stimulation at lower intensity which in turn lowers current requirement. This would decrease the draw on the batteries of implantable devices, which could extend the time between battery charges for patients with these devices, among other potential benefits. Future studies may assess the trade-off between efficacy and selectivity for various epidural stimulation paradigms and the optimization of epidurally implanted electrode arrays in humans.

## 5 Additional information

### 5.1 Competing interests

Jason B. Carmel is a Founder and stockholder in BackStop Neural and a scientific advisor and stockholder in SharperSense. Walter Voit co-founded several neuromodulation companies including Qualia Oto, Backstop Neural, and Kardiodia which have licenses to technology developed at UT Dallas. The other authors have nothing to disclose.

### 5.2 Funding

This work was supported by the National Institute of Neurological Disorders and Stroke, National Institutes of Health (award R01NS124224, 1R03NS141040), and a Department of Health New York State Institutional Grant (award C38336GG), and by the Congressionally Directed Medical Research Programs (award HT9425–24-1–0218).

## 5.3 Acknowledgements

We thank Jacob Kang for his assistance during the stimulation experiments. We thank Windsor Ting and Maria Bandres for their comments on the manuscript. We thank Columbia University’s Neurology High Performance Computing (HPC) Infrastructure and the Department of Systems Biology IT (NIH: S10OD032433, S10OD021764) for providing computational resources.

## 5.4 Author contributions

Conceptualization: ASA, JRM, JBC; Data Curation: VT, JRM; Formal Analysis: APL, VT, ASA, JRM; Funding Acquisition: WEV, JRM, JBC; Investigation: ASA; Methodology: ASA, PER, OR, WEV, JRM, JBC; Resources: PER, OR, WEV, JBC; Software: APL, VT, JRM; Supervision: WEV, JRM, JBC; Validation: APL, VT, ASA, JRM; Visualization: APL, VT, PER; Writing - Original Draft: APL, VT; Writing - Review & Editing: APL, VT, ASA, PER, OR, WEV, JRM, JBC.

## 5.5 Data availability

The datasets used in this study are publicly available [49].

## 5.6 Code availability

The code to reproduce the presented analyses is available at https://github.com/hbmep/rat-mapping.

## Supplementary information

### Methods

#### Surgical procedures

*Rats* Experiments were performed under anesthesia to enable testing with different devices. The rats were initially anesthetized with a mixture of ketamine (90 mg/kg) and xylazine (10 mg/kg), a combination commonly used in electrophysiological recordings [12, 38, 50]. Anesthesia levels were subsequently maintained using supplemental injection of ketamine at one-third of the initial dosage, administered as needed based on the continual assessment of the hindlimb pinch reflex. Following the initial anesthesia, the animals were placed in a stereotaxic frame with their heads fixed. Body temperature was maintained at 37.5 degrees during the procedure, with continuous monitoring using a rectal probe and heat delivered as needed by a warming pad. Blood oxygen levels were monitored with a pulse oximeter. Hair was removed from the top of the head, along the back, and over the right and left forelimbs. The skin was then cleaned with alcohol and an antiseptic solution. A midline incision was made, extending from between the eyes to the T1 vertebra. Connective tissue covering the skull was excised, and the muscles were separated to access the cervical spine. A hole was drilled into the skull over the olfactory bulbs and a screw electrode was inserted into the hole as a ground electrode.

##### EMG implant

EMG was recorded from eight different muscles: left extensor carpi radialis (ECR), flexor carpi radialis (FCR), biceps, triceps, abductor digiti quinti (ADQ), deltoid, biceps femoris, and right biceps. Flexible, braided stainless steel wires (AS631, Cooner Wire; Chatsworth, CA, USA) were employed for EMG recording. These electrodes were soldered to the connector and covered with epoxy to ensure insulation. Sub-sequently, the connector was attached to the recording system. Wire insulation was removed for 1 mm, starting from 2 cm away from the tip, corresponding to the muscle belly. A twenty-gauge needle was inserted into the muscles, and the wires were passed through the needle, allowing their tips to emerge on the opposite side of the muscle. The needle was then gently withdrawn while leaving the wire inside the muscle. Once the uninsulated part of the wire was positioned within the muscle belly, the wire tip was securely knotted and adhered to the muscle using gluture topical adhesive (World Precision Instruments; Florida, USA) to prevent potential movement. If necessary, the muscle was covered with skin and sutured to prevent it from drying out.

##### Spinal cord electrode placement

After placing the EMG electrodes, the muscles over the spinal cord were separated, and the T1 spinous process was clamped to stabilize the spine. A laminectomy was performed to bilaterally expose the C4 cervical spinal cord. Arrays were then carefully inserted into the dorsal epidural space over the cervical enlargement (C5-C8). We used bony and anatomical landmarks to identify the root entry zone. The intervertebral foramen also provided an opening for visual confirmation of the target site. Omnetics (Omnetics Connector Corp.; Minnesota, USA) connectors for the spinal array were attached to the stimulation system. Electrode impedance was checked both *in vitro* and *in vivo* before and after placement to ensure their quality.

### Results

**Supplementary Table S1.**
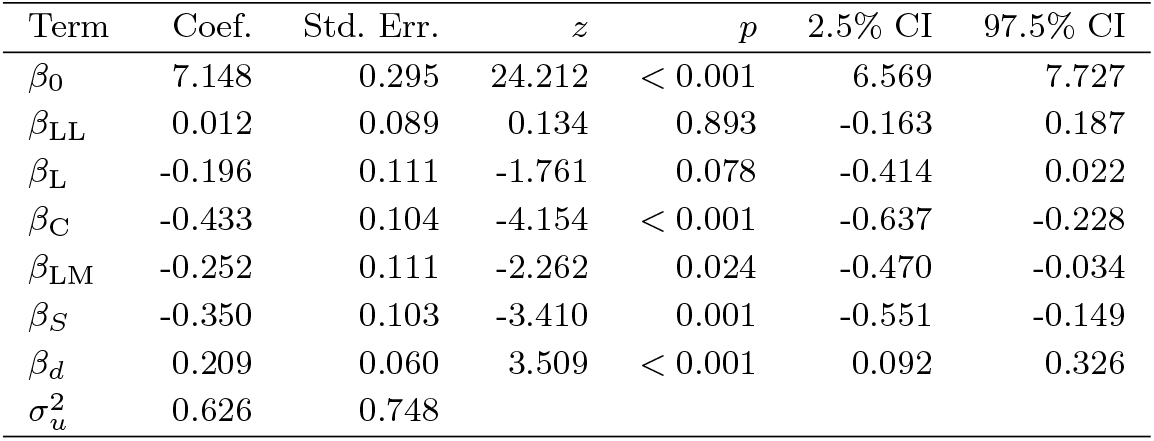
Mixed model output for effect of electrode position, distance, and size in linear array.

**Supplementary Table S2.** Mixed model output for effect of electrode position, distance, and orientation in circular array.

| Term | Coef. | Std. Err. | $z$ | $p$ | 2.5% CI | 97.5% CI |
| --- | --- | --- | --- | --- | --- | --- |
| $\beta_0$ | 6.955 | 0.393 | 17.711 | < 0.001 | 6.186 | 7.725 |
| $\beta_C$ | -0.798 | 0.153 | -5.207 | < 0.001 | -1.098 | -0.498 |
| $\beta_{NE}$ | -0.292 | 0.124 | -2.347 | 0.019 | -0.535 | -0.048 |
| $\beta_N$ | -0.822 | 0.137 | -5.978 | < 0.001 | -1.091 | -0.552 |
| $\beta_{NW}$ | -0.439 | 0.124 | -3.528 | < 0.001 | -0.682 | -0.195 |
| $\beta_W$ | -0.103 | 0.124 | -0.825 | 0.409 | -0.346 | 0.141 |
| $\beta_{SW}$ | -0.575 | 0.124 | -4.629 | < 0.001 | -0.819 | -0.332 |
| $\beta_S$ | -1.133 | 0.137 | -8.242 | < 0.001 | -1.402 | -0.864 |
| $\beta_{SE}$ | -0.478 | 0.124 | -3.841 | < 0.001 | -0.721 | -0.234 |
| $\beta_c$ | -0.206 | 0.090 | -2.294 | 0.022 | -0.382 | -0.030 |
| $\beta_s$ | -0.053 | 0.092 | -0.573 | 0.567 | -0.232 | 0.127 |
| $\beta_d$ | 0.481 | 0.141 | 3.418 | 0.001 | 0.205 | 0.757 |
| $\sigma_u^2$ | 1.156 | 1.732 | | | | |

**Supplementary Table S3.**
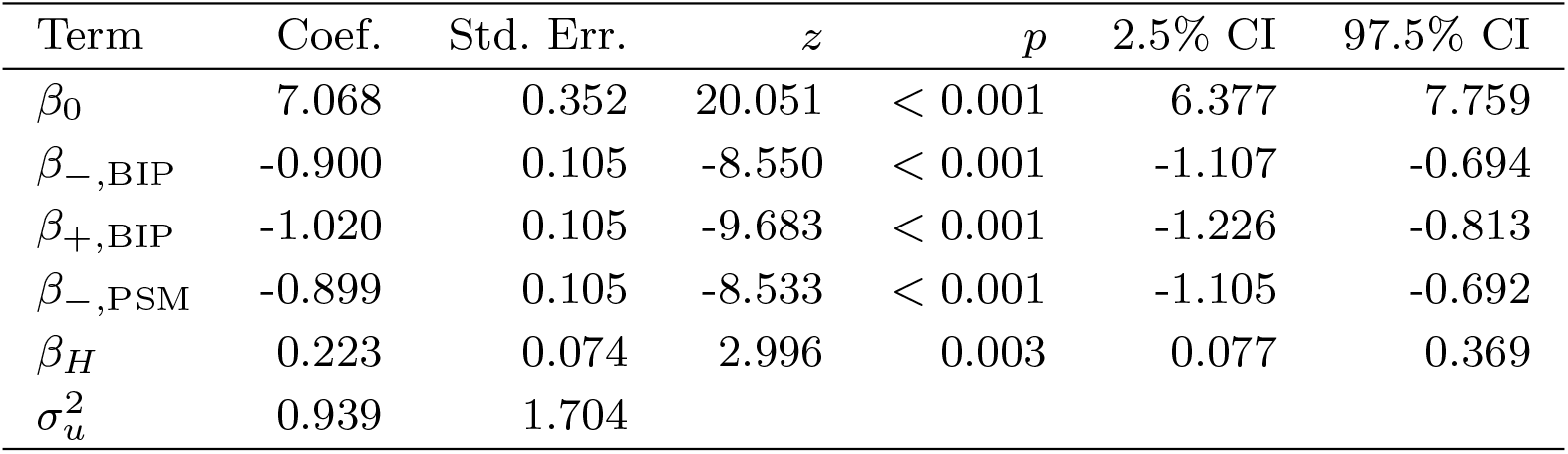
Mixed model output for effect of polarity, waveform, and high-definition in circular array.

## Notes

### Summary of Updates

We replaced pairwise comparisons with unified linear mixed effects models to isolate and compare relative impact of key parameters including electrode distance, orientation, size, and position.

## References

[1] Rocha-Flores, P.E., Chitrakar, C., Rodriguez-Lopez, O., Ren, Y., Joshi-Imre, A., Parikh, A.R., Asan, A.S., McIntosh, J.R., Garcia-Sandoval, A., Pancrazio, J.J., Ecker, M., Lu, H., Carmel, J.B., Voit, W.E.: Softening, Conformable, and Stretchable Conductors for Implantable Bioelectronics Interfaces. Advanced Materials Technologies 10(6), 2401047 (2025) 10.1002/admt.202401047

[2] Garcia-Sandoval, A., Guerrero, E., Hosseini, S.M., Rocha-Flores, P.E., Rihani, R., Black, B.J., Pal, A., Carmel, J.B., Pancrazio, J.J., Voit, W.E.: Stable softening bioelectronics: A paradigm for chronically viable ester-free neural interfaces such as spinal cord stimulation implants. Biomaterials 277, 121073 (2021) 10.1016/j.biomaterials.2021.121073

[3] Garcia-Sandoval, A., Pal, A., Mishra, A.M., Sherman, S., Parikh, A.R., Joshi-Imre, A., Arreaga-Salas, D., Gutierrez-Heredia, G., Duran-Martinez, A.C., Nathan, J., Hosseini, S.M., Carmel, J.B., Voit, W.: Chronic softening spinal cord stimulation arrays. Journal of Neural Engineering 15(4), 045002 (2018) 10.1088/1741-2552/aab90d

[4] Rattay, F., Minassian, K., Dimitrijevic, M.R.: Epidural electrical stimulation of posterior structures of the human lumbosacral cord: 2. quantitative analysis by computer modeling. Spinal Cord 38(8), 473–489 (2000) 10.1038/sj.sc.3101039

[5] Capogrosso, M., Wenger, N., Raspopovic, S., Musienko, P., Beauparlant, J., Luciani, L.B., Courtine, G., Micera, S.: A Computational Model for Epidural Electrical Stimulation of Spinal Sensorimotor Circuits. Journal of Neuroscience 33(49), 19326–19340 (2013) 10.1523/JNEUROSCI.1688-13.2013

[6] Greiner, N., Barra, B., Schiavone, G., Lorach, H., James, N., Conti, S., Kaeser, M., Fallegger, F., Borgognon, S., Lacour, S., Bloch, J., Courtine, G., Capogrosso, M.: Recruitment of upper-limb motoneurons with epidural electrical stimulation of the cervical spinal cord. Nature Communications 12(1), 435 (2021) 10.1038/s41467-020-20703-1

[7] Renshaw, B.: Activity in the simplest spinal reflex pathways. Journal of Neurophysiology 3(5), 373–387 (1940) 10.1152/jn.1940.3.5.373

[8] Lloyd, D.P.C.: Reflex action in relation to pattern and peripheral source of afferent stimulation. Journal of Neurophysiology 6(2), 111–119 (1943) 10.1152/jn.1943.6.2.111

[9] Yang, Q., Ramamurthy, A., Lall, S., Santos, J., Ratnadurai-Giridharan, S., Zareen, N., Alexander, H., Ryan, D., Martin, J.H., Carmel, J.B.: Independent replication of motor cortex and cervical spinal cord electrical stimulation to promote forelimb motor function after spinal cord injury in rats. Experimental Neurology 320, 112962 (2019) 10.1016/j.expneurol.2019.112962

[10] Pal, A., Park, H., Ramamurthy, A., Asan, A.S., Bethea, T., Johnkutty, M., Carmel, J.B.: Spinal cord associative plasticity improves forelimb sensorimotor function after cervical injury. Brain 145(12), 4531–4544 (2022) 10.1093/brain/awac235

[11] Rattay, F., Danner, S.M., Hofstoetter, U.S., Minassian, K.: Finite element modeling for extracellular stimulation. In: Jaeger, D., Jung, R. (eds.) Encyclopedia of Computational Neuroscience, pp. 1–12. Springer, New York, NY (2014). 10.1007/978-1-4614-7320-6593-5

[12] Mishra, A.M., Pal, A., Gupta, D., Carmel, J.B.: Paired motor cortex and cervical epidural electrical stimulation timed to converge in the spinal cord promotes lasting increases in motor responses. The Journal of Physiology 595(22), 6953–6968 (2017) 10.1113/JP274663

[13] Wenger, N., Moraud, E.M., Gandar, J., Musienko, P., Capogrosso, M., Baud, L., Le Goff, C.G., Barraud, Q., Pavlova, N., Dominici, N., Minev, I.R., Asboth, L., Hirsch, A., Duis, S., Kreider, J., Mortera, A., Haverbeck, O., Kraus, S., Schmitz, F., DiGiovanna, J., Van Den Brand, R., Bloch, J., Detemple, P., Lacour, S.P., Bézard, E., Micera, S., Courtine, G.: Spatiotemporal neuromodulation therapies engaging muscle synergies improve motor control after spinal cord injury. Nature Medicine 22(2), 138–145 (2016) 10.1038/nm.4025

[14] McIntosh, J.R., Joiner, E.F., Goldberg, J.L., Murray, L.M., Yasin, B., Mendiratta, A., Karceski, S.C., Thuet, E., Modik, O., Shelkov, E., Lombardi, J.M., Sardar, Z.M., Lehman, R.A., Mandigo, C., Riew, K.D., Harel, N.Y., Virk, M.S., Carmel, J.B.: Intraoperative electrical stimulation of the human dorsal spinal cord reveals a map of arm and hand muscle responses. Journal of Neurophysiology 129(1), 66–82 (2023) 10.1152/jn.00235.2022

[15] Zannou, A.L., Koochesfahani, M.B., Gaugain, G., Nikolayev, D., Russo, M., Bikson, M.: Computational optimization of spinal cord stimulation for dorsal horn interneuron polarization. Neuromodulation: Technology at the Neural Interface 28(6), 952–961 (2025) 10.1016/j.neurom.2025.01.015

[16] Holsheimer, J., Struijk, J.J., Tas, N.R.: Effects of electrode geometry and combination on nerve fibre selectivity in spinal cord stimulation. Medical and Biological Engineering and Computing 33(5), 676–682 (1995) 10.1007/BF02510785

[17] Zander, H.J., Kowalski, K.E., DiMarco, A.F., Lempka, S.F.: Model-Based Optimization of Spinal Cord Stimulation for Inspiratory Muscle Activation. Neuromodulation: Journal of the International Neuromodulation Society 25(8), 1317–1329 (2022) 10.1111/ner.13415

[18] North, R.B., Kidd, D.H., Olin, J.C., Sieracki, J.M.: Spinal cord stimulation electrode design: prospective, randomized, controlled trial comparing percutaneous and laminectomy electrodes-part I: technical outcomes. Neurosurgery 51(2), 381–389389390 (2002) 10.1097/00006123-200208000-00015

[19] North, R.B., Kidd, D.H., Petrucci, L., Dorsi, M.J.: Spinal cord stimulation electrode design: a prospective, randomized, controlled trial comparing percutaneous with laminectomy electrodes: part II-clinical outcomes. Neurosurgery 57(5), 990–996990996 (2005) 10.1227/01.neu.0000180030.00167.b9

[20] Mao, G.-W., Zhang, J.-J., Su, H., Zhou, Z.-J., Zhu, L.-S., Lü, X.-Y., Wang, Z.-G.: A flexible electrode array for determining regions of motor function activated by epidural spinal cord stimulation in rats with spinal cord injury. Neural Regeneration Research 17(3), 601–607 (2022) 10.4103/1673-5374.320987

[21] Russman, S.M., Montgomery-Walsh, R., Vatsyayan, R., U, H.S., Diaz-Aguilar, L.D., Chang, E.Y., Tang, Q., Lee, K., Yaksh, T.L., Ben-Haim, S., Ciacci, J., Dayeh, S.A.: The Conformal, High-Density SpineWrap Microelectrode Array for Focal Stimulation and Selective Muscle Recruitment. Advanced Functional Materials 35(16), 2420488 (2025) 10.1002/adfm.202420488

[22] Struijk, J.J., Holsheimer, J., Boom, H.B.: Excitation of dorsal root fibers in spinal cord stimulation: a theoretical study. IEEE transactions on bio-medical engineering 40(7), 632–639 (1993) 10.1109/10.237693

[23] Coburn, B.: A theoretical study of epidural electrical stimulation of the spinal cord–Part II: Effects on long myelinated fibers. IEEE transactions on bio-medical engineering 32(11), 978–986 (1985) 10.1109/TBME.1985.325649

[24] Alam, M., Garcia-Alias, G., Shah, P.K., Gerasimenko, Y., Zhong, H., Roy, R.R., Edgerton, V.R.: Evaluation of optimal electrode configurations for epidural spinal cord stimulation in cervical spinal cord injured rats. Journal of Neuroscience Methods 247, 50–57 (2015) 10.1016/j.jneumeth.2015.03.012

[25] Alam, M., Garcia-Alias, G., Jin, B., Keyes, J., Zhong, H., Roy, R.R., Gerasimenko, Y., Lu, D.C., Edgerton, V.R.: Electrical neuromodulation of the cervical spinal cord facilitates forelimb skilled function recovery in spinal cord injured rats. Experimental Neurology 291, 141–150 (2017) 10.1016/j.expneurol.2017.02.006

[26] Ladenbauer, J., Minassian, K., Hofstoetter, U.S., Dimitrijevic, M.R., Rattay, F.: Stimulation of the human lumbar spinal cord with implanted and surface electrodes: a computer simulation study. IEEE transactions on neural systems and rehabilitation engineering: a publication of the IEEE Engineering in Medicine and Biology Society 18(6), 637–645 (2010) 10.1109/TNSRE.2010.2054112

[27] Zander, H.J., Graham, R.D., Anaya, C.J., Lempka, S.F.: Anatomical and technical factors affecting the neural response to epidural spinal cord stimulation. Journal of Neural Engineering 17(3), 036019 (2020) 10.1088/1741-2552/ab8fc4

[28] Cuellar, C., Lehto, L., Islam, R., Mangia, S., Michaeli, S., Lavrov, I.: Selective Activation of the Spinal Cord with Epidural Electrical Stimulation. Brain Sciences 14(7), 650 (2024) 10.3390/brainsci14070650

[29] Rattay, F.: Analysis of Models for External Stimulation of Axons. IEEE Transactions on Biomedical Engineering BME-33(10), 974–977 (1986) 10.1109/TBME.1986.325670

[30] Meacham, K.W., Giuly, R.J., Guo, L., Hochman, S., DeWeerth, S.P.: A lithographically-patterned, elastic multi-electrode array for surface stimulation of the spinal cord. Biomedical Microdevices 10(2), 259–269 (2008) 10.1007/s10544-007-9132-9

[31] Nandra, M.S., Lavrov, I.A., Edgerton, V.R., Tai, Y.-C.: A parylene-based microelectrode array implant for spinal cord stimulation in rats. In: 2011 IEEE 24th International Conference on Micro Electro Mechanical Systems, pp. 1007–1010 (2011). 10.1109/MEMSYS.2011.5734598

[32] Minev, I.R., Musienko, P., Hirsch, A., Barraud, Q., Wenger, N., Moraud, E.M., Gandar, J., Capogrosso, M., Milekovic, T., Asboth, L., Torres, R.F., Vachicouras, N., Liu, Q., Pavlova, N., Duis, S., Larmagnac, A., Vörös, J., Micera, S., Suo, Z., Courtine, G., Lacour, S.P.: Electronic dura mater for long-term multimodal neural interfaces. Science 347(6218), 159–163 (2015) 10.1126/science.1260318

[33] Rocha-Flores, P.E., Guerrero, E., Rodríguez-Lopez, O., Chitrakar, C., Parikh, A.R., Pancrazio, J.J., Cogan, S.F., Ecker, M., Voit, W.E.: Softening and flexible hybrid electronics integration for biomedical applications. MRS Communications 13(5), 892–900 (2023) 10.1557/s43579-023-00431-5

[34] Datta, A., Elwassif, M., Battaglia, F., Bikson, M.: Transcranial current stimulation focality using disc and ring electrode configurations: FEM analysis. Journal of Neural Engineering 5(2), 163–174 (2008) 10.1088/1741-2560/5/2/007

[35] Datta, A., Bansal, V., Diaz, J., Patel, J., Reato, D., Bikson, M.: Gyri-precise head model of transcranial direct current stimulation: improved spatial focality using a ring electrode versus conventional rectangular pad. Brain Stimulation 2(4), 201–2072071 (2009) 10.1016/j.brs.2009.03.005

[36] Tyagi, V., Murray, L.M., Asan, A.S., Mandigo, C., Virk, M.S., Harel, N.Y., Carmel, J.B., McIntosh, J.R.: Hierarchical Bayesian estimation of motor-evoked potential recruitment curves yields accurate and robust estimates. Brain Stimulation 18(6), 1855–1870 (2025) 10.1016/j.brs.2025.09.008

[37] Cogan, S.F.: Neural Stimulation and Recording Electrodes. Annual Review of Biomedical Engineering 10(1), 275–309 (2008) 10.1146/annurev.bioeng.10.061807.160518. Accessed 2023-06-11

[38] Sharma, P., Shah, P.K.: In vivo electrophysiological mechanisms underlying cervical epidural stimulation in adult rats. The Journal of Physiology 599(12), 3121–3150 (2021) 10.1113/JP281146

[39] Seabold, S., Perktold, J.: Statsmodels: Econometric and statistical modeling with python. SciPy 2010 (2010) 10.25080/Majora-92bf1922-011

[40] Holm, S.: A Simple Sequentially Rejective Multiple Test Procedure. Scandinavian Journal of Statistics 6(2), 65–70 (1979)

[41] Faria, P., Hallett, M., Miranda, P.C.: A finite element analysis of the effect of electrode area and inter-electrode distance on the spatial distribution of the current density in tDCS. Journal of Neural Engineering 8(6), 066017 (2011) 10.1088/1741-2560/8/6/066017. Accessed 2026-04-02

[42] Troughton, J.G., Ansong Snr, Y.O., Duobaite, N., Proctor, C.M.: Finite element analysis of electric field distribution during direct current stimulation of the spinal cord: Implications for device design. APL Bioengineering 7(4), 046109 (2023) 10.1063/5.0163264

[43] Janssen, A.M., Oostendorp, T.F., Stegeman, D.F.: The coil orientation dependency of the electric field induced by TMS for M1 and other brain areas. Journal of NeuroEngineering and Rehabilitation 12(1), 47 (2015) 10.1186/s12984-015-0036-2

[44] Rawji, V., Ciocca, M., Zacharia, A., Soares, D., Truong, D., Bikson, M., Rothwell, J., Bestmann, S.: tDCS changes in motor excitability are specific to orientation of current flow. Brain Stimulation 11(2), 289–298 (2018) 10.1016/j.brs.2017.11.001

[45] Foerster, A., Yavari, F., Farnad, L., Jamil, A., Paulus, W., Nitsche, M.A., Kuo, M.-F.: Effects of electrode angle-orientation on the impact of transcranial direct current stimulation on motor cortex excitability. Brain Stimulation 12(2), 263–266 (2019) 10.1016/j.brs.2018.10.014

[46] Reijonen, J., Säisänen, L., Könönen, M., Mohammadi, A., Julkunen, P.: The effect of coil placement and orientation on the assessment of focal excitability in motor mapping with navigated transcranial magnetic stimulation. Journal of Neuroscience Methods 331, 108521 (2020) 10.1016/j.jneumeth.2019.108521

[47] Coburn, B., Sin, W.K.: A theoretical study of epidural electrical stimulation of the spinal cord–Part I: Finite element analysis of stimulus fields. IEEE transactions on bio-medical engineering 32(11), 971–977 (1985) 10.1109/tbme.1985.325648

[48] Geddes, L.A., Baker, L.E.: The specific resistance of biological material—A compendium of data for the biomedical engineer and physiologist. Medical and biological engineering 5(3), 271–293 (1967) 10.1007/BF02474537

[49] Pascual-Leone, A., Tyagi, V., Asan, A., Rocha-Flores, P.E., Rodriguez Lopez, O., Voit, W., McIntosh, J.R., Carmel, J.: Dataset: Electrode Position, Size, and Orientation Determine Efficacy of Cervical Epidural Stimulation to Recruit Forelimb Muscles in Rats. 10.5281/zenodo.17107079

[50] Ahmed, Z.: Dipolar cortico-muscular electrical stimulation: a novel method that enhances motor function in both - normal and spinal cord injured mice. Journal of NeuroEngineering and Rehabilitation 7(1), 46 (2010) 10.1186/1743-0003-7-46

